# Cognitive reserve counteracts typical neural activity changes related to ageing

**DOI:** 10.1101/2023.02.01.526583

**Authors:** Jesús Cespón, Irina Chupina, Manuel Carreiras

**Affiliations:** BCBL Basque Center on Cognition, Brain, and Language, Donostia/San Sebastián, Spain; Radboud University, Donders Centre for Cognition, Nijmegen, The Netherlands; Ikerbasque. Basque Foundation for Science, Bilbao, Spain; University of the Basque Country (UPV/EHU). Bilbao, Spain

**Keywords:** brain maintenance, cognitive reserve, event-related potentials, executive functions, neurocognitive ageing theories

## Abstract

Studies have shown that older adults with high Cognitive Reserve (HCR) exhibit better executive functioning than their low CR (LCR) counterparts. However, the neural processes linked to those differences are unclear. This study investigates (1) the neural processes underlying enhanced executive functions in older adults with HCR and (2) how executive control differences are modulated by task difficulty. We recruited 74 participants, who performed two executive control tasks with different difficulty levels while recording the electroencephalogram. The accuracy on both tasks requiring inhibition of irrelevant information was better in the HCR than the LCR group. Also, in the more demanding task, event-related potentials (ERP) latencies related to inhibition and working memory update were faster in HCR than LCR. Moreover, the HCR, but not the LCR, showed larger P300 amplitude in parietal than frontal regions and in the left than right hemisphere, suggesting a posterior to anterior shift of activity and loss of inter-hemispheric asymmetries in LCR participants. These results suggest that high CR counteracts neural activity changes related to ageing. Thus, high levels of CR can be related to maintenance of neural activity patterns typically observed in young adults rather than to deployment of neural compensatory mechanisms.

## 1. Introduction

The construct of cognitive reserve (CR) was proposed to explain why some individuals are more resistant than others to brain pathology (Katzman et al., 1988; Stern, 2012). As stated by Stern (2012), older adults with high CR exhibit enhanced cognitive functioning and delayed age of dementia diagnosis than older adults with low CR. High levels of CR can be developed through exposure to cognitively stimulating experiences (e.g., high educational level and occupational status), which are frequently used as proxy variables to estimate CR levels (Chapko et al., 2018). Although there is considerable evidence for CR from epidemiological studies, the neural bases of CR are still far from clear (Steffener and Stern, 2012). A better understanding of the neural activity patterns linked to CR would be useful to detect individuals at risk of cognitive impairment and to investigate to what extent interventions as well as specific variables and lifestyle factors (e.g., multilingualism, playing musical instruments, and physical exercise) contribute to induce high CR neural activity patterns.

Considering that ageing has been associated with impaired executive functions (Ferguson et al., 2021), which are a set of cognitive processes (e.g., inhibition, attentional switching, and working memory update) that are crucial to carrying out daily life activities (e.g., driving, cooking, shopping), several studies have investigated whether high levels of CR could counteract such executive impairment in older adults. In this context, several studies reported that older adults with high CR exhibit better performance at tasks tapping executive functions compared to their low CR counterparts (Corral et al., 2006; Darby et al., 2017; Oosterman et al., 2021; Roldán-Tapia et al., 2012).

To clarify the neural correlates of CR, some studies have used event-related brain potentials (ERP) to investigate neural processing in older adults with different levels of CR during the performance of executive tasks (Gu et al., 2018; Quinzi et al., 2020; Speer and Soldan, 2015). These studies, which focused mainly on the P300 ERP –a correlate of working memory update (Polich, 2007) – have produced inconsistent results. For instance, Quinzi et al (2020) showed larger P300 amplitude in high than low CR. However, other studies did not find such differences (Gu et al., 2018; Speer and Soldan, 2015). Similarly, Speer and Soldan (2015) reported faster P300 latency in high than low CR but later research using different task paradigms did not find differences (Gu et al., 2018; Quinzi et al., 2020). To the best of our knowledge, electrophysiological studies did not focus on whether and how global neural activity patterns related to ageing –namely, posterior to anterior shift of activity (Davis et al., 2008) and inter-hemispheric dedifferentiation (Cabeza, 2002) - are modulated by CR. Studying this issue would be important to shed light on the neural activity patterns related to successful cognitive ageing (McDonough et al., 2022). To date, some studies have linked these neural activity patterns to compensatory mechanisms that preserve cognition (Cabeza, 2002; Davis et al., 2008) but others proposed that preserved cognition is associated with maintenance of young like neural activity patterns (e.g., Koen and Rugg, 2019; Morcom and Henson, 2018).

A recent review carried out by Balart-Sánchez et al (2021) emphasizes that, even if electrophysiological measures are sensitive to CR, the results are largely influenced by the task design. In fact, studying correlates of CR requires taking into account the difficulty of the task since behavioural and/or neural differences related to CR may emerge only at high task difficulty levels (Martinez et al., 2022). In addition, many previous studies have used small samples, which diminish the reliability and replicability of the results (Button et al., 2013).

In order to study neural correlates of CR, we recruited a sample of 74 older adults, who performed two spatial stimulus-response compatibility (SRC) tasks with different difficulty level; namely, the Simon task and the spatial Stroop task. Both these tasks allow studying inhibition (as participants have to inhibit the tendency to react towards the attended location) and attentional switching (as participants have to switch and update the stimulus-response binding on a trial to trial basis) skills (for details about the tasks, see methods). However, the spatial Stroop is more difficult than the Simon task due to the higher number of conflicting information sources (Cespón et al., 2020).

ERP studies of CR using spatial SRC tasks have mainly analysed frontal N200 and parietal P300 (Cespón et al., 2020). P300 is a correlate of stimulus-response binding emerging from parietal regions and is delayed and attenuated in incongruent compared to congruent trials (Cespón et al., 2020). Frontal N200, an ERP linked to inhibition and conflict monitoring (Folstein and Van Petten, 2008), increases in incongruent compared to congruent trials in young adults but such differences are often blurred at advanced stages of life (Cespón and Carreiras, 2020), probably as a consequence of decreased signal/noise ratio and/or lower synchronisation among neural systems (Gajewski et al., 2018). A few ERP studies have also focused on the central contralateral negativity component (N2cc), which was linked to premotor activity to prevent the tendency to react towards the attended location (Praamstra and Oostenveld, 2003). Here, we analysed N200, P300, and N2cc, as they each detect key aspects of executive processing during spatial SRC tasks.

The objectives of the present study are to: 1) obtain neural correlates of CR during the performance of executive tasks; 2) investigate to what extent such correlates are modulated by task difficulty; 3) study how posterior-to-anterior shift of activity and inter-hemispheric dedifferentiation are modulated by CR in order to assess to what extent high levels of CR relate to deployment of neural compensatory mechanisms or maintenance of neural activity patterns observed in young adults.

For the first objective, we hypothesized that the high CR (HCR) group would show faster ERP latencies and increased ERP amplitudes compared to the low CR (LCR) group, reflecting neural differences related to ageing (Cespón and Carreiras, 2020; Gajewski et al., 2018). Regarding the second objective, we predicted that such ERP differences would occur only (or would be stronger) in the more demanding task (i.e., the spatial Stroop task). For the third objective, we expected that, in comparison to HCR group, the LCR group would show a larger posterior-to-anterior shift of activity within the P300 time window and increased loss of hemispheric asymmetry, as revealed by the absence of ERP amplitude differences between left and right frontal and parietal regions of interest within N200 and P300 time windows.

## 2. Methods

### 2.1. Participants

74 older adults (age range: 61-82 years old) took part in this research, although data from 5 participants was discarded due to EEG artefacts. CR was estimated by using a standardized questionnaire (Rami et al., 2011). Following previous studies (Quinzi et al., 2020) we divided participants into low vs. high CR groups using the median CR score as a cut-off point. 35 participants (mean age: 69.6; standard deviation (SD): 4.9) were assigned to the HCR group (mean CR score: 18.1; SD: 2.21) and 34 participants (mean age: 72.2; SD: 4.3) were assigned to the LCR group (mean CR score: 11.6; SD: 2.64). Participants were right-handed, as assessed by the Edinburgh handedness inventory test (Oldfield, 1971), and had normal or corrected to normal visual acuity. Participants reported no previous history of neurological or psychiatric disorders. Before starting the experiment, participants were informed about the procedures of the study and provided written informed consent to take part in the research. This research was performed in compliance with the ethical guidelines defined by the Declaration of Helsinki and received prior approval from the local Ethics Committee.

### 2.2. Experimental procedures and tasks

All the participants took part in a first experimental session, which involved a general neuropsychological assessment. Specifically, participants performed the Mini-mental state examination (Folstein, 1975), the Spanish version of the Repeatable Battery for the Assessment of Neuropsychological Status (De la Torre et al., 2004; Randolph, 1998), and the CR questionnaire (Rami et al., 2011). All participants performed at a level that indicated preserved cognitive functioning. Participants took part in a second experimental session to perform two executive control tasks (i.e., a Simon task and a spatial Stroop task) during EEG recording.

The participants performed Simon and spatial Stroop tasks (see Figure 1). In the Simon task, participants responded to the colour of a lateralised square (by pressing a left or a right response button with the corresponding hand) while ignoring its location (the square appeared in the right or the left hemifield). In the spatial Stroop task, participants responded according to the arrow direction (pointing to the right or to the left) while ignoring its location (the arrow appeared in the right or the left hemifield). For both tasks, participants were instructed to direct their gaze to the centre of the screen, where a central fixation cross appeared for 500ms against a black background. Then, the lateralized stimulus (i.e., a square in the Simon task and an arrow in the spatial Stroop task) appeared for 100ms. To prevent exogenous activity in the electroencephalogram (EEG), a non-target stimulus of similar shape and lateralised location was simultaneously presented in the contralateral hemifield with respect to the target stimulus. The target and non-target stimuli were presented 5cm far from the central fixation cross. Participants sat 100cm in front of the computer screen and the entire display was presented within the foveal region (Bargh and Chartrand, 2000). After stimulus presentation, the screen remained blank for a period of 2000±250ms. Next, a new trial started with the appearance of the central fixation cross. In each task, participants were instructed to respond as fast and accurately as possible to the target stimulus. In the spatial Stroop task, a stimulus-stimulus conflict (e.g., a left lateralised arrow pointing to the right, see Figure 1) covaries with the stimulus-response conflict and for this reason it is thought to be more difficult than the Simon task (Juncos-Rabadán et al., 2008; Lu and Proctor, 1995). For each experimental task, 120 trials per condition were presented, giving rise to a total of 480 trials (i.e., 120 × 4 conditions: congruent-Congruent (c-C), incongruent-Congruent (i-C), incongruent-Incongruent (i-I), and congruent-Incongruent (c-I). Each task was divided into three blocks of 160 trials each. Participants performed a practice block of 10 trials before starting each task and rested for about one minute between blocks.

**Figure 1.**
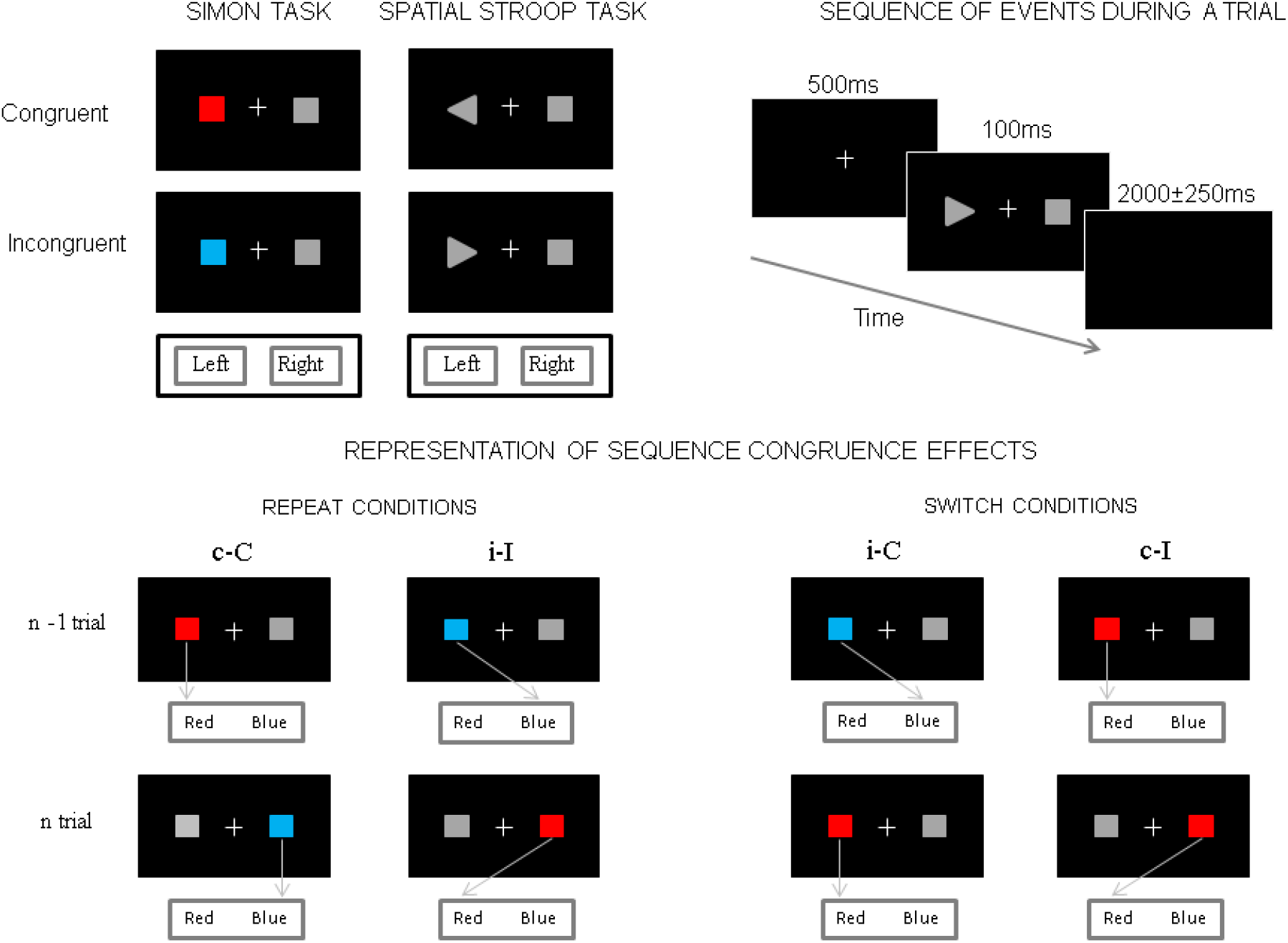
Representation of the experimental tasks and conditions. Left top panel: congruent and incongruent conditions for the Simon task and the spatial Stroop task; Right top panel: sequence of events during a trial (identical parameters were set for both tasks). Bottom panel: experimental conditions that were analysed: congruent-congruent (c-C), incongruent-congruent (i-C), incongruent-incongruent (i-I), and congruent-Incongruent (c-I), being the first member of the pairs (lowercase letters) the “n-1” trial and the second member of the pairs (capital letters) the “n” trial. Note that, in switching conditions, the stimulus-response binding has to be changed from the “n-1” to “n” trial.

### 2.3. EEG recording

The continuous EEG was recorded using Easycap (Brain Products GmbH, Germany). Fifty-four EEG electrodes were placed on the scalp, according to the international 5-10 system for positioning. Vertical and horizontal electrooculogram signals were recorded by two electrodes located above and below the right eye and two electrodes located in the outer part of the lateral canthus of both eyes, respectively, in order to correct artefacts related to ocular movements. The ground electrode was placed at Fpz. The right mastoid served as an online reference for all electrodes, whereas the left mastoid electrode was used offline to re-reference the scalp recordings to the average of the left and the right mastoid, i.e., including the implicit reference (right mastoid) in the calculation of the new reference. The EEG signal was acquired with a bandpass filter of 0.01-1000 Hz and digitized at a sampling rate of 1000 Hz. The impedance values were kept below 5 kΩ for all electrodes.

### 2.4. Data analyses

Behavioural performance was evaluated by analysing RTs and accuracy (i.e., number of errors -NE). RTs slower than 1500 ms were automatically excluded from all analyses. ERPs were calculated for correct responses only. The EEG signal was filtered with a 0.1-80 Hz digital bandpass and a 50 Hz notch filter. Ocular and muscular artefacts were eliminated through independent component analysis [algorithm Infomax (Gradient) in Brain Vision Analyzer 2.2]. Epochs exceeding ±100 μV were automatically rejected and those still displaying artefacts were manually removed from subsequent analysis. The epochs were established between −200 and 1000ms relative to the onset of the target stimulus. N200 latency was identified as the maximum negative peak between 200ms and 400ms after the stimulus presentation. P300 latency was identified as the maximum positive peak between 300ms and 600ms after the stimulus presentation. N200 and P300 amplitudes were analysed using the mean amplitude in time windows of 50 and 100ms, respectively. P300 was analysed in midline electrodes (Fz, Cz, Pz), which also allowed us to investigate the posterior-to-anterior shift of activity. The frontal N200 was analysed at Fz, as this was where it showed its maximum amplitude.

To compare brain activity between both hemispheres, we pooled the following electrodes to create left and right regions of interest (ROIs): left frontal (F3, F5, FC3, FC5), right frontal (F4, F6, FC4, FC6), left parietal (P3, P5, CP3, CP5), and right parietal (P4, P6, CP4, CP6). N200 amplitude was studied in frontal ROIs (as we were interested in frontal N200, which is related to executive functions). P300 amplitude was studied in frontal and parietal ROIs because these regions have been related to attentional control and working memory update processes, respectively, within the time range of P300 (Polich, 2007). In order to compute N200 and P300 amplitudes we followed the same procedures as described for midline electrodes.

In order to obtain indexes of frontalisation, we carried out the subtraction “Fz – Pz” for P300 amplitudes (higher values correspond to greater frontalisation). We focused on midline sites since it is where P300 reaches maximum amplitude. To study dedifferentiation, we carried out the subtractions “Right ROI – Left ROI” in frontal and parietal ROIs for P300 amplitudes and in frontal ROIs for N200 amplitudes. Activity is left lateralised in spatial SRC tasks (Cespón et al., 2020; Spironelli et al., 2006). So, higher dedifferentiation is indicated by larger values for P300 and lower values for N200.

In line with previous studies (Amenedo et al., 2012; Cespón et al., 2022), N2cc was calculated based on the hemifield of the target stimulus location by applying the following formula: [C4 - C3 (left hemifield stimuli) + C3 - C4 (right hemifield stimuli)] / 2]. We obtained the N2cc waveform regardless of whether the stimulus location was congruent or incongruent with the required response. This procedure does not allow comparison between conditions but it has the advantage that residual motor activity is removed from the N2cc waveform (Cespón et al., 2020). Specifically, half of the stimuli located in the left hemifield require a left-handed response, whereas the other half requires a right-handed response. Averaging across all these trials, motor activity is cancelled out. Importantly, as the target stimulus is always located in the left hemifield, target-related activity remains. The same reasoning can be applied to averages for right-hemifield stimuli. The N2cc peak latency was identified as the largest negative peak between 200–400ms after stimulus presentation. For each participant, the N2cc amplitude was calculated in a time window of 50ms (i.e., ±25ms around peak latency).

### 2.5. Statistical analyses

We carried out repeated measures ANOVAs with Task (two levels: Simon task, spatial Stroop task), Congruence (two levels: Congruent, Incongruent), and Switching (two levels: Repeat, Switch) as within-subject factors and CR (two levels: HCR, LCR) as between-subjects factor for RT, NE, and N200 latency and amplitude. For P300 latency and amplitude we additionally included Electrode as a within-subject factor (three levels: Fz, Cz, and Pz). For N2cc we carried out a repeated measures ANOVA with Task (two levels: Simon task, spatial Stroop task) as the within-subject factor and CR (two levels: HCR, LCR) as the between-subjects factor. We carried out repeated measures ANOVAs for frontal (N200 and P300 amplitudes) and parietal (P300 amplitudes) ROIs using Task (two levels: Simon task, spatial Stroop task), Congruence (two levels: Congruent, Incongruent), Switching (two levels: Repeat, Switch), and Hemisphere (two levels: Left, Right) as within-subject factors and CR (two levels: HCR, LCR) as a between-subjects factor. Also, we conducted independent samples t-tests to study differences between LCR and HCR in the frontalisation and dedifferentiation indexes.

When ANOVAs showed significant effects related to the main factors and/or their interactions, pairwise comparisons were conducted using Bonferroni correction. We applied the Greenhouse-Geisser correction for degrees of freedom if the condition of sphericity was not met. Partial eta square (η2p), which is an effect size measure, has been provided for significant results.

## 3. Results

### 3.1. Behavioural results

The repeated measures ANOVA (Group x Task x Congruence x Sequence) for RTs (see Table 1, top part) showed an effect of Task [F (1, 67) = 28.7, p < 0.001, η^2^p = 0.300], with longer RTs in the spatial Stroop (i.e., the more demanding task) than in the Simon task (p < 0.001). There were also significant effects of Congruence [F (1, 67) = 215.5, p < 0.001, η^2^p = 0.763], with slower RTs in incongruent than congruent trials (p < 0.001) and sequence [F (1, 67) = 314.5, p < 0.001, η^2^p = 0.824], as RTs were slower in switch than repeat trials (p < 0.001).

**Table 1.** Reaction times (RT) and Number of Errors (NE) for each group -low cognitive reserve (LCR) and high cognitive reserve (HCR)- and experimental condition –congruent-congruent (c-C), incongruent-Congruent (i-C), incongruent-incongruent (i-I), and congruent-Incongruent (c-I) in the Simon and spatial Stroop tasks. Significant differences are described in the main text.

|  | **SIMON TASK** |  |  |  | **SPATIAL STROOP TASK** |  |  |  |
| --- | --- | --- | --- | --- | --- | --- | --- | --- |
|  | **REACTION TIMES** |  |  |  | **REACTION TIMES** |  |  |  |
|  | c-C | i-C | i-I | c-I | c-C | i-C | i-I | c-I |
| **LCR** | 550.3<br>(72.2) | 577.9<br>(82.5) | 605.5<br>(87.9) | 622.1<br>(84.5) | 583.6<br>(73.8) | 629.8<br>(90.5) | 677.9<br>(114.5) | 704.1<br>(123.8) |
| **HCR** | 534.5<br>(60.8) | 561.6<br>(66.1) | 582.4<br>(57.6) | 605.1<br>(68.2) | 554.3<br>(82.9) | 590.3<br>(86.8) | 628.7<br>(79.7) | 660.0<br>(101.0) |
|  | **NUMBER OF ERRORS** |  |  |  | **NUMBER OF ERRORS** |  |  |  |
| **LCR** | 1.17<br>(1.58) | 3.29<br>(3.56) | 7.32<br>(6.62) | 10.41<br>(7.17) | 1.41<br>(1.90) | 2.64<br>(3.09) | 6.29<br>(5.63) | 10.26<br>(7.80) |
| **HCR** | 0.88<br>(1.38) | 2.00<br>(2.48) | 3.37<br>(3.07) | 6.14<br>(4.54) | 0.65<br>(1.62) | 1.57<br>(2.35) | 4.05<br>(4.05) | 6.08<br>(4.91) |

The repeated measures ANOVA (Group x Task x Congruence x Sequence) for NE (see Table 1, bottom part) showed a significant Group effect [F (1, 67) = 9.44 p = 0.003, η^2^p = 0.124], as the NE was higher in the LCR compared to the HCR group (p = 0.003). There were also significant effects of congruence [F (1, 67) = 105.3, p < 0.001, η^2^p = 0.611], as the NE was higher in incongruent than congruent trials (p < 0.001) and sequence [F (1, 67) = 135.7, p < 0.001, η^2^p = 0.670], as the NE was higher in switch than repeat trials (p < 0.001). The Group × Congruence interaction reached significance [F (1, 67) = 8.16, p = 0.006, η^2^p = 0.109], as the NE in the incongruent condition was higher in the LCR compared to the HCR group (p = 0.003), whereas there was no significant difference in the congruent condition.

### 3.2. Event-related brain potentials

#### 3.2.1. Analyses in the midline electrodes

The repeated measures ANOVA (Group × Task × Congruence × Sequence) for N200 latency (Figure 2) showed an effect of Congruence [F (1, 67) = 4.39, p = 0.040, η^2^p = 0.062], as N200 was longer in incongruent than congruent trials (p < 0.001). For N200 amplitude, the analysis showed a significant effect of Task [F (1, 67) = 4.57, p = 0.036, η^2^p = 0.064], as N200 was larger in the spatial Stroop than in the Simon task (p = 0.036) and Congruence [F (1, 67) = 23.8, p < 0.001, η^2^p = 0.262], as N200 was larger in incongruent than congruent trials (p < 0.001).

**Figure 2.**
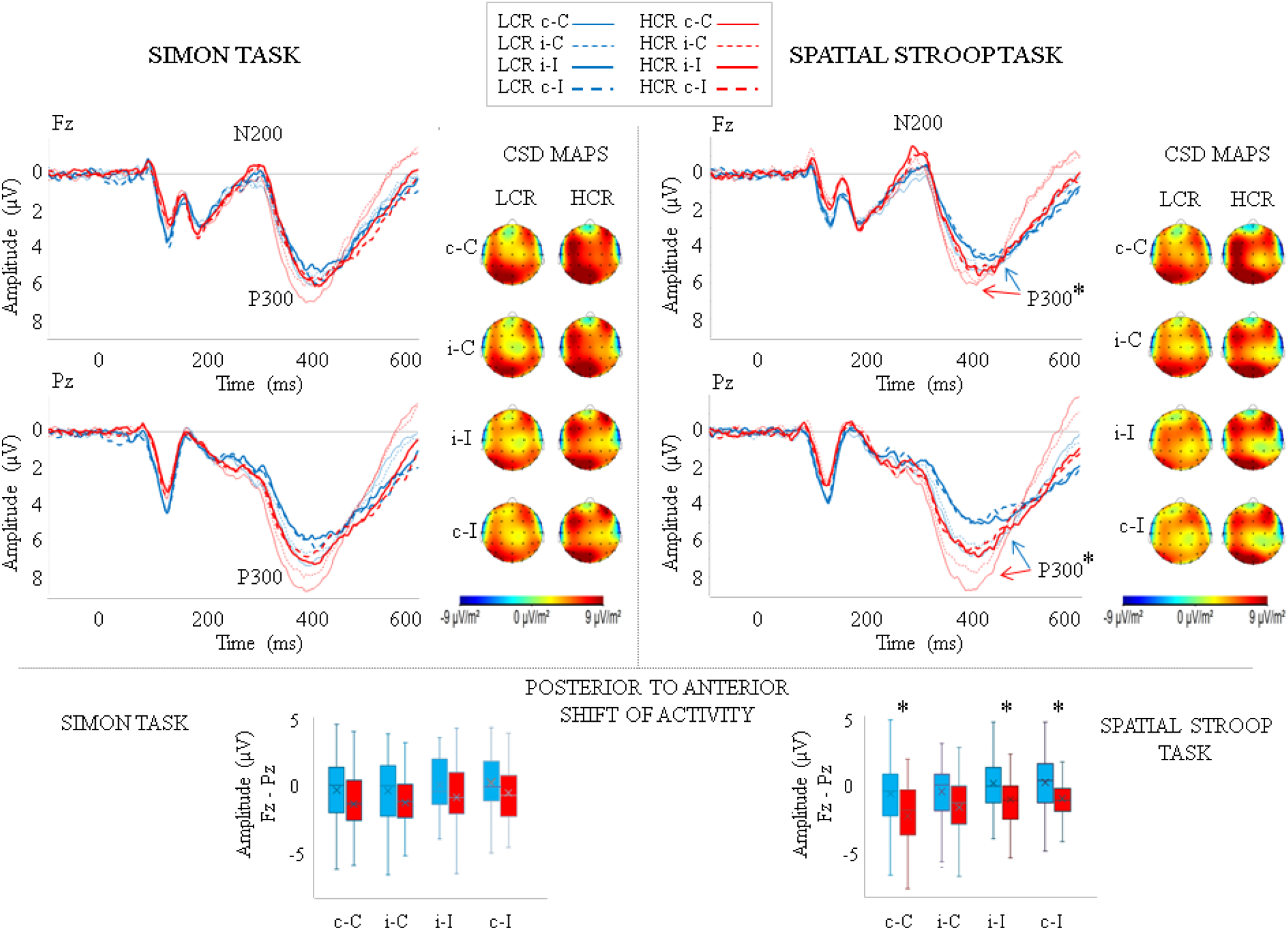
The top part of the figure shows the waveforms in the four conditions of the Simon and spatial Stroop tasks at Fz and Pz as well as the current density (CSD) maps for low cognitive reserve (LCR) and high cognitive reserve (HCR) groups. CSD maps were made by taking a 100ms time window around P300 peak latency for each experimental condition. In the spatial Stroop task, the P300 peak latency was longer in LCR than HCR. The bottom part of the figure shows a box and whiskers chart for Simon and spatial Stroop tasks with the frontalisation indexes on each condition. Frontalisation was higher in LCR than HCR in c-C and i-I conditions of the spatial Stroop task.

The repeated measures ANOVA (Group × Task × Congruence × Sequence × Electrode) for P300 latency (Figure 2) showed a significant Task × Group interaction [F (1, 67) = 9.56, p = 0.003, η^2^p = 0.125]. Specifically, in the spatial Stroop task the P300 was slower in LCR compared to HCR group (p < 0.001) whereas in the Simon task no differences were observed. Also, a Task effect was observed [F (1, 67) = 12.8, p = 0.001, η^2^p = 0.161], as P300 was slower in the spatial Stroop than in the Simon task (p = 0.001). A Congruence effect was also observed for P300 latency [F (1, 67) = 30.4, p < 0.001, η^2^p = 0.312], as P300 was slower in incongruent than congruent trials (p < 0.001). For P300 amplitude, the repeated measures ANOVA showed an Electrode x Group interaction effect [F (2, 134) = 3.75, p = 0.026, η^2^p = 0.053]. In HCR, P300 was larger in Cz than Fz (p < 0.001) as well as in Pz than Fz (p < 0.001). In LCR, P300 was larger in Cz than Fz (p = 0.026) but differences between Pz and Fz were not significant. Also, a significant effect of Task was observed [F (1, 67) = 6.31, p = 0.014, η^2^p = 0.086]. Namely, P300 was larger in the Simon than in the spatial Stroop task (p = 0.014). An effect of Congruence was observed [F (1, 67) = 29.8, p < 0.001, η^2^p = 0.308], as P300 was larger in congruent than incongruent trials (p < 0.001). The effect of Sequence was significant [F (1, 67) = 18.2, p < 0.001, η^2^p = 0.214], as P300 was larger in repeat than switch trials (p < 0.001). For the frontalisation indexes (see Figure 2, lower panel), independent sample t-tests revealed increased frontalisation in LCR than HCR in c-C (t (67) = 1.42, p = 0.018), i-I (t (67) = 2.28, p = 0.026), and c-I (t (67) = 2.23, p = 0.029) conditions of the spatial Stroop task but no differences in the Simon task.

#### 3.2.2. Negativity central contralateral (N2cc)

The repeated measures ANOVA (Group x Task) for N2cc latency (Figure 3) showed a Task effect [F (1, 67) = 47.9, p < 0.001, η^2^p = 0.417], as the N2cc was slower in the spatial Stroop than in the Simon task (p < 0.001). The Group effect was significant [F (1, 67) = 4.94, p = 0.030, η^2^p = 0.069], as N2cc was slower in LCR than HCR (p = 0.030). For N2cc amplitude, the repeated measures ANOVA showed a Task effect [F (1, 67) = 54.7, p < 0.001, η^2^p = 0.450], as N2cc was larger in spatial Stroop than Simon task. The Task × Group interaction was marginally significant [F (1, 67) = 3.83, p = 0.054, η^2^p = 0.054] as N2cc was larger in HCR than LCR only in the spatial Stroop task (p = 0.051).

**Figure 3.**
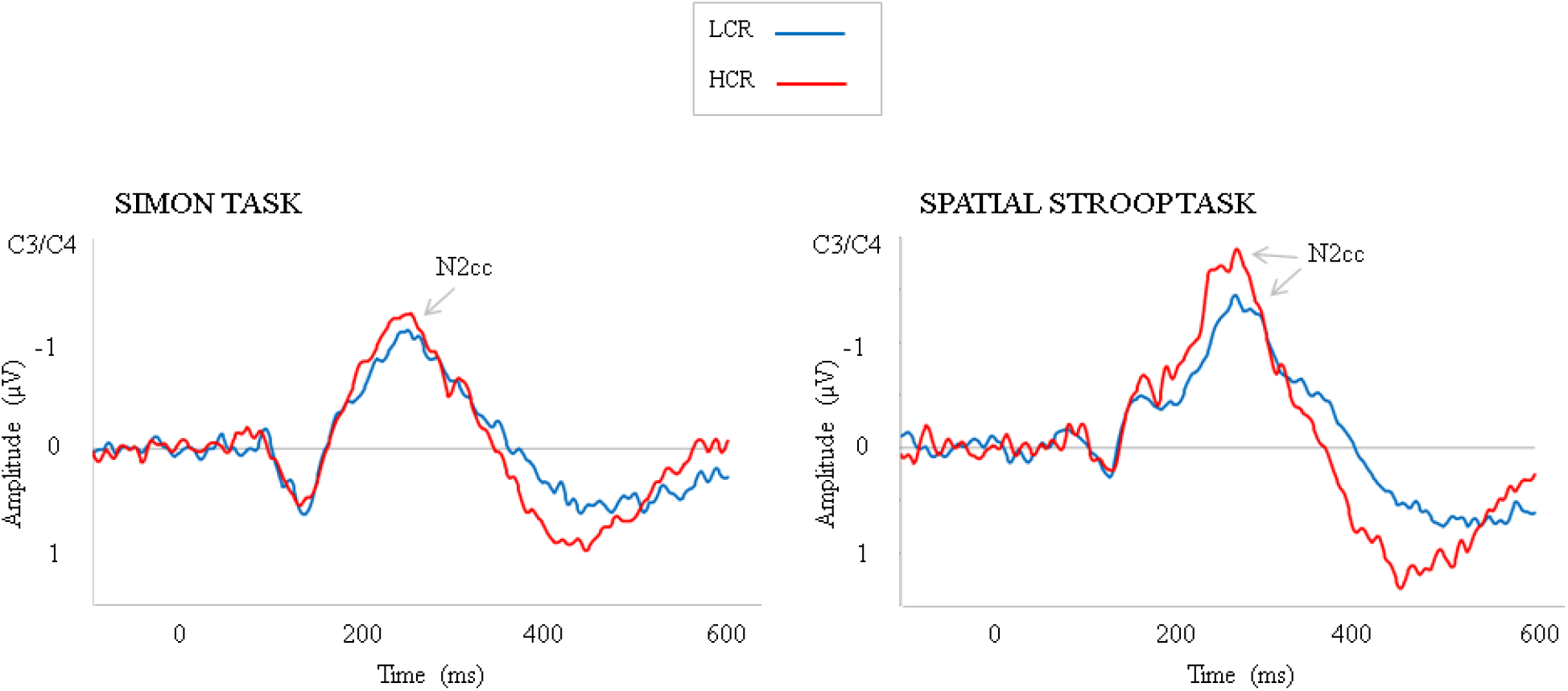
Negativity central contralateral (N2cc). The amplitude of N2cc was larger and its latency slower in the spatial Stroop than in the Simon task. For the spatial Stroop task, N2cc peak latency was slower in LCR than HCR group and N2cc amplitude was larger in HCR compared to LCR group.

#### 3.2.3. Analyses in regions of interest

The repeated measures ANOVA (Group x Task x Congruence x Sequence x Hemisphere) for N200 amplitude (Figure 4) showed a Congruence effect [F (1, 67) = 22.95, p < 0.001, η^2^p = 0.255], as N200 was larger in incongruent than congruent trials (p < 0.001). The analysis showed a ROI effect [F (1, 67) = 13.40, p < 0.001, η^2^p = 0.167], as N200 was larger in the left than in the right hemisphere (p < 0.001). The repeated measures ANOVA (Group x Task x Congruence x Sequence x Hemisphere) for frontal P300 amplitude (Figure 4) showed a ROI x Group interaction [F (1, 67) = 5.66, p < 0.020, η^2^p = 0.078], as the HCR group showed larger frontal P300 in left than right hemisphere (p < 0.001) but differences were not significant in the LCR group. A Task effect was significant [F (1, 67) = 7.49, p = 0.008, η^2^p = 0.101], as frontal P300 was larger in Simon than spatial Stroop task (p = 0.008). The Congruence effect was significant [F (1, 67) = 13.05, p = 0.001, η^2^p = 0.163], as P300 was larger in congruent than incongruent trials (p = 0.001). A Sequence effect was shown [F (1, 67) = 8.55, p = 0.005, η^2^p = 0.113], as frontal P300 was larger in repeat than switch trials (p = 0.005). For dedifferentiation indexes in frontal ROIs (Figure 4, right side charts), analyses for frontal P300 revealed that differentiation was larger in LCR than HCR in c-C (t (67) = 2.62, p = 0.011) and i-I (t (67) = 2.40, p = 0.019) conditions of the spatial Stroop task but there were no differences in the Simon task. No differences were found in the N200 time window.

**Figure 4.**
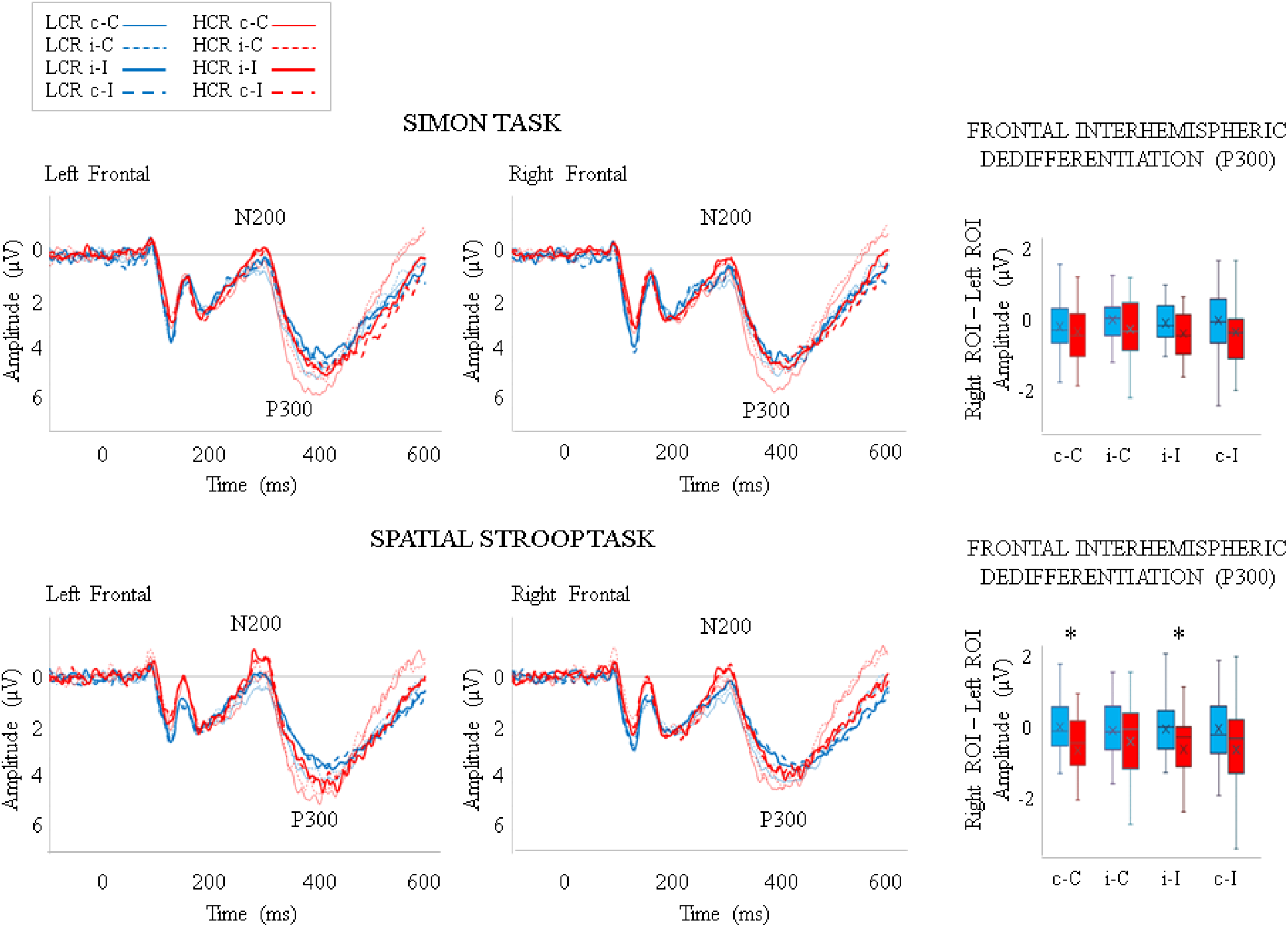
Event-related potentials in left and right frontal regions of interest (ROI). Frontal P300 amplitude was larger in left than right frontal region for HCR group. No differences were found for the LCR group. On the right side of the figure, dedifferentiation indexes are represented for both groups through a box and whiskers chart. Dedifferentiation was higher in LCR than HCR group in the spatial Stroop task (significant differences are represented by an asterisk).

The repeated measures ANOVA (Group × Task × Congruence × Sequence × Hemisphere) for parietal P300 amplitude (Figure 5) showed a ROI × Group interaction effect [F (1, 67) = 4.88, p = 0.031, η^2^p = 0.068]; namely, in the HCR, parietal P300 was larger in the left than in the right hemisphere (p < 0.001) but such differences were not observed in the LCR group. Also, this analysis revealed a Task effect [F (1, 67) = 7.94, p = 0.006, η^2^p = 0.106], as the parietal P300 was larger in the Simon than in the spatial Stroop task (p = 0.006). Moreover, the analysis revealed an effect of Congruence [F (1, 67) = 47.6, p < 0.001, η^2^p = 0.416], as the parietal P300 was larger in congruent than incongruent trials (p = 0.006). Sequence showed a significant effect [F (1, 67) = 13.3, p < 0.001, η^2^p = 0.224], as P300 was larger in repeat than switch trials (p < 0.001). The dedifferentiation indexes in parietal ROIs (Figure 5, right side charts) revealed higher dedifferentiation in LCR than HCR in c-C (t (67) = 2.077, p = 0.042), i-C (t (67) = 2.02, p = 0.047), and i-I (t (67) = 2.56, p = 0.012) conditions of the spatial Stroop task and i-I condition (t (67) = 2.02, p = 0.047) of the Simon task.

**Figure 5.**
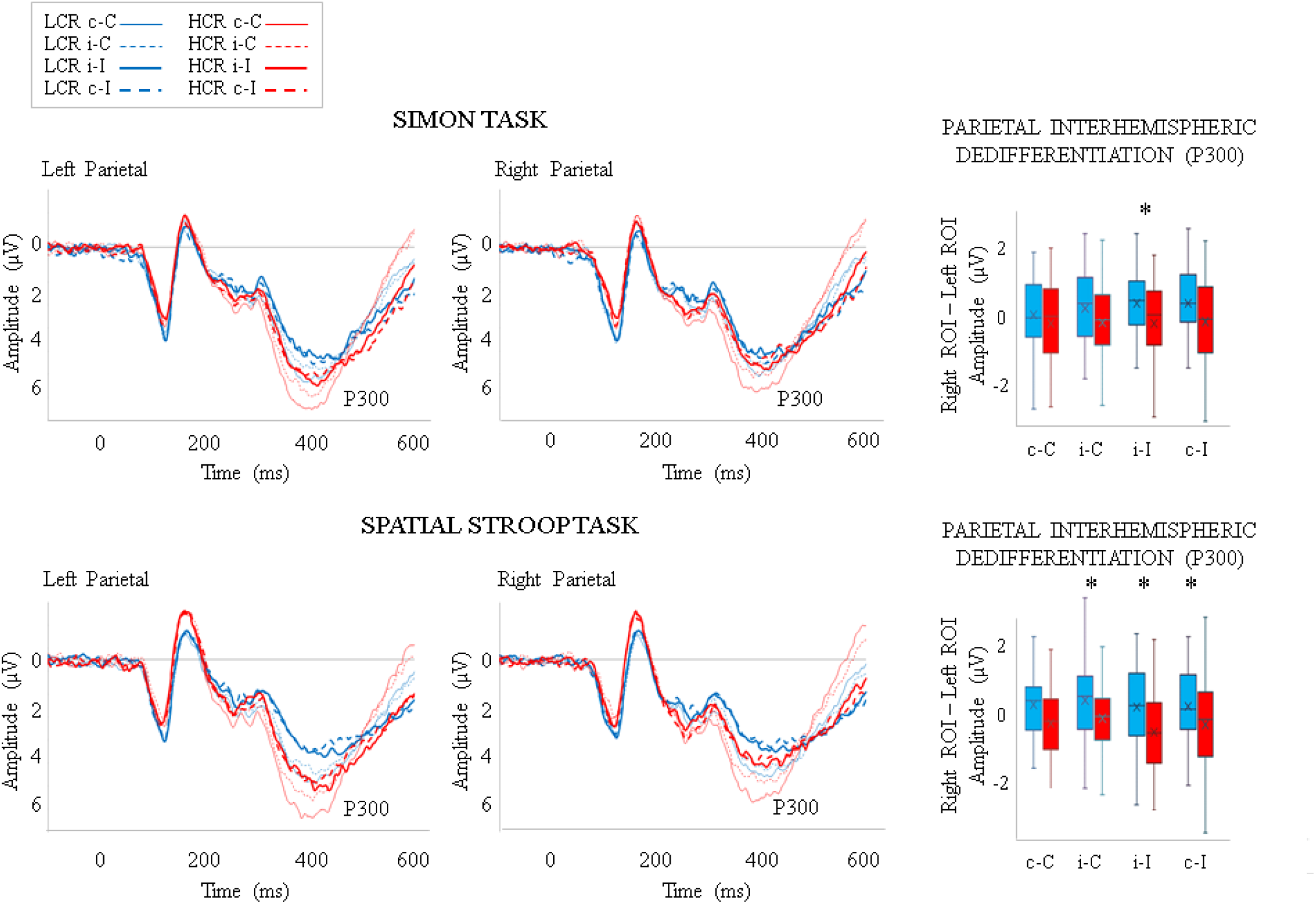
Event-related potentials in left and right parietal regions of interest (ROI). Parietal P300 amplitude was larger in the left than in the right parietal region for the HCR group but no differences were found for the LCR group. On the right side of the figure, dedifferentiation indexes are represented for both groups through a box and whiskers chart. Dedifferentiation was higher in LCR than HCR group for both tasks (significant differences are represented by an asterisk).

## 4. Discussion

The results of the present study showed greater accuracy in the HCR compared to the LCR group in incongruent conditions on both tasks, indicating better executive functions in HCR than LCR, although RTs did not differ between both groups. P300 and N2cc latencies were faster in HCR than LCR group in the spatial Stroop task but not in the Simon task. The HCR group also showed larger P300 amplitudes in parietal than frontal electrodes and larger P300 amplitudes in left than right frontal and parietal ROIs but such differences were not observed in the LCR group, suggesting a posterior to anterior shift of activity and loss of inter-hemispheric asymmetries, respectively.

In line with previous findings, RTs were slower when the stimulus location was incongruent (i.e., in i-I and c-I trials) than congruent (i.e., in c-C and i-C trials) with the side of the required response (Cespón et al., 2020; Lu and Proctor, 1995). This interference effect reflects additional processing time to inhibit the response based on the stimulus location (Cespón et al., 2020; Valle-Inclán, 1996). In addition, RTs were longer in switch (i.e., i-C and c-I) compared to repeat (i.e., c-C and i-I) conditions, which shows the existence of sequence congruence effects, as previously reported by studies using SRC tasks (Lamers and Roelofs, 2011; Spapè and Hommel, 2014). The increased RT in the switch compared to repeat condition may be interpreted as the time required to switch and update the S-R binding that was active in working memory in the previous trial (Cespón et al., 2020). The present research also showed that RTs were slower on the spatial Stroop compared to the Simon task, supporting the assumption that the spatial Stroop task is more challenging than the Simon task (Lu and Proctor, 1995; Juncos-Rabadán et al., 2008). This increased difficulty of the spatial Stroop task compared to the Simon task allowed us to study the effect of task difficulty on the existence of differences between HCR and LCR groups.

The present study showed better executive skills in high than low CR older adults, which were revealed by higher response accuracy in the HCR compared to LCR group. These results align with previous studies reporting enhanced executive functions in high CR than low CR older adults (Corral et al., 2006; Darby et al., 2017; Oosterman et al., 2021; Roldán-Tapia et al., 2012). Importantly, accuracy was higher in HCR than LCR in the spatially incongruent conditions of both tasks, but not in switching conditions. This suggests that the HCR group exhibited better inhibition skills compared to the LCR group rather than a general advantage in information processing speed or a specific advantage in switching and update of working memory contents. However, sequence congruence effects in older adults may be largely modulated by the inter-trial interval (Aisenberg et al., 2014). Therefore, further research studying differences between low and high CR at different inter-trial intervals is needed before strong conclusions can be drawn about the impact of CR on switching skills related to sequence congruence effects.

The present research revealed that high CR is associated with faster ERP latencies and enhanced ERP amplitudes, which suggests that older adults with high CR exhibit more youthful neural activity patterns compared to low CR older adults. The faster N2cc latency in the HCR than the LCR group indicates that the allocation of inhibitory activity related to preventing the bias to respond towards the spatial location (Praamstra and Oostenveld, 2003; Praamstra, 2006; Cespón et al., 2020) was faster in HCR than LCR group in the spatial Stroop task. Thus, age-related slowing in deploying neural processes related to the N2cc (Amenedo et al., 2012; Cespón et al., 2022) can be counteracted to some extent by high levels of CR. Moreover, faster P300 latency in the HCR than LCR group suggested faster updating of the S-R binding in HCR than LCR in the four conditions of the spatial Stroop task (Cespón et al., 2020; Hoppe et al., 2017), showing that high CR reduces the age-related slowing in neural processes related to P300 that has been reported by previous studies (Van der Lubbe and Verleger, 2002; Zurrón et al., 2014). For ERP amplitudes, N2cc was larger in HCR than LCR in the spatial Stroop task but such differences were not observed in the Simon task. As ERP amplitudes are usually reduced with physiological ageing (Cespón and Carreiras, 2020; Gajewski et al., 2018), these results are in line with the hypothesized relationship between high CR and younger neural activity patterns.

Evidence for the increased difficulty of the spatial Stroop compared to the Simon task was also provided by the ERPs analysed. Specifically, frontal N200 amplitude was larger in the spatial Stroop than in the Simon task, which aligns with increased frontal N200 amplitudes at increased difficulty level (Folstein and Van Petten, 2008). P300 amplitude was also larger in the Simon than in the spatial Stroop task, which is consistent with theoretical assumptions that P300 amplitude decreases as task difficulty level increases (Polich, 2007). Moreover, N2cc latency was slower and N2cc amplitude was larger in the spatial Stroop than in the Simon task, which reflects slowed and additional neural activity to prevent the response towards the stimulus location in the spatial Stroop task (Cespón et al., 2020).

The results of the present study show the importance of task difficulty in the appearance of neural differences between HCR and LCR groups because such differences (namely, faster N2cc and P300 latencies and larger N2cc amplitude in HCR than LCR) only appeared in the more challenging task (i.e., the spatial Stroop task). These results are in line with previous behavioural (Martinez et al., 2022) and ERP (Speer and Soldan, 2015) studies showing differences between low and high CR groups only at higher task difficulty levels. Even so, some recent studies focusing on P300 modulations by CR failed to find differences between high and low CR groups in the more demanding task (e.g., Gu et al., 2018). Importantly, Gu et al. (2018) found that, P300 differences between low and high demanding tasks were larger in the low than in the high CR group. This pattern of results suggests that differences in P300 could have been found in Gu et al (2018) if the difficulty level of the more challenging task had been higher.

For the global neural activity patterns, the HCR group showed larger P300 amplitude in parietal than frontal sites but such differences were not observed in LCR group, which may be interpreted as a frontalisation of neural activity in the LCR group. This interpretation is supported by the frontalisation indices in the spatial Stroop task (i.e., the more difficult task) and it is consistent with recent studies (Kuruvilla-Mathew et al., 2022; Morcom and Henson, 2018). Also, the left and right frontal and parietal ROIs showed increased activity in the left compared to right hemisphere for the HCR group in the P300 time window but not for the LCR group. These results point to the loss of inter-hemispheric asymmetries in low CR older adults during the performance of spatial SRC tasks, in which left lateralised hemisphere activity in younger adults has been shown (Spironelli et al., 2006). This interpretation is supported by the dedifferentiation indexes analysed in both spatial SRC tasks. Thus, the results of the present study align with studies showing negative relationships between increased inter-hemispheric dedifferentiation and cognitive functioning in older adults (Knights et al., 2021; Koen and Rugg, 2019; Morcom and Jonson, 2015). Finally, differences between HCR and LCR groups in inter-hemispheric asymmetries for P300 but not for N200 are consistent with a recent ERP study (Tagliabue et al., 2022), suggesting that reduced inter-hemispheric asymmetrical activity associated with ageing does not uniformly affect all stages of information processing.

The findings of the present study (that is, faster ERP latencies and larger ERP amplitudes in the HCR than LCR group in addition to higher activity in the left than right hemisphere as well as in parietal than frontal regions in the HCR but not in the LCR group) suggest that high CR is related to preservation of brain activity patterns observed in young adults rather than to deployment of compensatory mechanisms. According to the scaffolding theory of aging and cognition (Reuter-Lorenz and Park, 2014), positive lifetime experiences counteract the brain insults related to ageing. In line with results from interventions based on cognitive training and physical exercise, it is possible that compensatory neural activity (i.e., increased neural activity associated with enhanced performance) is manifested when a new skill or positive lifestyle activity is being introduced but such brain activity trend to be reduced after that new skill or habit is consolidated (Cespón et al., 2018). Taking into account that habits, lifestyles, and variables related to high CR in older adults were probably established several decades ago, the benefits related to variables associated with high CR are manifested by preserved brain activity patterns rather than deployment of compensatory mechanisms.

## 5. Conclusions

The present study showed that inhibitory skills were better in older adults with high CR than low CR, as revealed by group-related differences in accuracy in the spatially incongruent conditions. In addition, faster N2cc and P300 latencies and increased N2cc amplitudes were evidenced in HCR than LCR in the more challenging task, suggesting an important role of task difficulty in the appearance of neural differences between HCR and LCR groups. Also, the LCR group (but not the HCR group) showed brain activity patterns typically associated with the aged brain; namely, posterior to anterior shift of activity and loss of inter-hemispheric asymmetry. The obtained findings suggest that HCR levels can counteract neural activity changes related to ageing. Thus, high levels of CR are associated with the maintenance of neural activity patterns observed in young adults (i.e., faster latencies, enhanced amplitudes, and differentiated brain activity patterns) rather than with the deployment of neural compensatory mechanisms. In response to requirements formulated in recent studies (Cespón, 2021; de Bruin et al., 2021), the results of the present study provide an important experimental foundation to interpret whether specific variables and lifestyle factors (such as physical exercise, playing music, or speaking two or more languages) increase executive functions and CR by inducing neural activity patterns associated with high CR.

## Acknowledgments

This project has received funding from the European Union’s Horizon 2020 research and innovation programme under the Marie Skłodowska-Curie grant agreement No. 838536, Spanish Ministry of Science (PID2019-105538RA-I00), from the Basque Government through the BERC 2022-2025 program, and from the Agencia Estatal de Investigación through BCBL’s Severo Ochoa excellence award CEX2020-001010-S.

The funders have/had no role in study design, data collection and analysis, decision to publish or preparation of the manuscript.

## Authors Contributions

Conceptualization: JC, MC; Data curation: JC, IC; Formal analysis: JC, IC; funding acquisition: JC, MC; Methodology: JC, MC; Supervision: MC; Writing original draft: JC; Review & editing: MC.

